# MERITS: a web-based integrated *Mycobacterial* PE/PPE protein database

**DOI:** 10.1101/2023.12.26.573374

**Authors:** Zhijie He, Cong Wang, Xudong Guo, Heyun Sun, Yue Bi, Miranda E. Pitt, Chen Li, Jiangning Song, Lachlan JM Coin, Fuyi Li

## Abstract

**Motivation:** PE/PPE proteins, highly abundant in the *Mycobacterium* genome, play a vital role in virulence and immune modulation. Understanding their functions is key to comprehending the internal mechanisms of *Mycobacterium*. However, a lack of dedicated resources has limited research into PE/PPE proteins.

**Results:** Addressing this gap, we introduce MERITS, a comprehensive 3D structure database specifically designed for PE/PPE proteins. MERITS hosts 22,353 non-redundant PE/PPE proteins, encompassing details like physicochemical properties, subcellular localisation, post-translational modification sites, protein functions, and measures of antigenicity, toxicity, and allergenicity. MERITS also includes data on their secondary and tertiary structure, along with other relevant biological information. MERITS is designed to be user-friendly, offering interactive search and data browsing Features to aid researchers in exploring the potential functions of PE/PPE proteins. MERITS is expected to become a crucial resource in the field, aiding in developing new diagnostics and vaccines by elucidating the sequence-structure-functional relationships of PE/PPE proteins.

**Availability and implementation:** MERITS is freely accessible at http://merits.unimelb-biotools.cloud.edu.au/.

## 1 Introduction

The *Mycobacterium* genus, characterized by its elongated, slightly curved, and occasionally branched bacilli, is widespread in both external environments and within human and animal hosts. Notably, pathogenic species such as *M. tuberculosis* and *M. leprae* pose significant health risks, causing severe diseases like tuberculosis and leprosy that result in respiratory, cutaneous, and mucosal infections (Johansen *et al*., 2020; Saxena *et al*., 2021; Sampson, 2011). These diseases underline the urgent need for effective interventions. Central to this challenge is the unique PE/PPE protein families found in *Mycobacteria*, named after their conserved proline (P) and glutamic acid (E) residues (Wang *et al*., 2020; Chandra *et al*., 2022). These proteins play critical roles in immune evasion, virulence, and pathogenicity, making them key to understanding and combating *Mycobacterial* diseases. Detailed analysis of the sequences, structures, and functions of PE/PPE proteins is crucial for developing new therapeutic strategies (Ehtram *et al*., 2021; Li *et al*., 2023).

Currently, researchers rely on general databases like NCBI (O’Leary *et al*., 2016), UniProt (Consortium *et al*., 2023), RCSB PDB (Kouranov, 2006), and AlphaFold DB (Varadi *et al*., 2022) for studying PE/PPE proteins. While these databases provide valuable genomic, transcript, and protein sequence records, their coverage of PE/PPE proteins is limited, offering only basic sequence information and lacking comprehensive annotation and detailed tertiary structures. Although UniProt, RCSB PDB, and AlphaFold DB focus on protein data, they fall short in providing extensive data on PE/PPE proteins. This lack of a dedicated database for PE/PPE proteins hinders in-depth research and limits our understanding of their role in *Mycobacterial* pathogenesis. Consequently, the development of a specialized database becomes imperative.

**Fig. 1.**
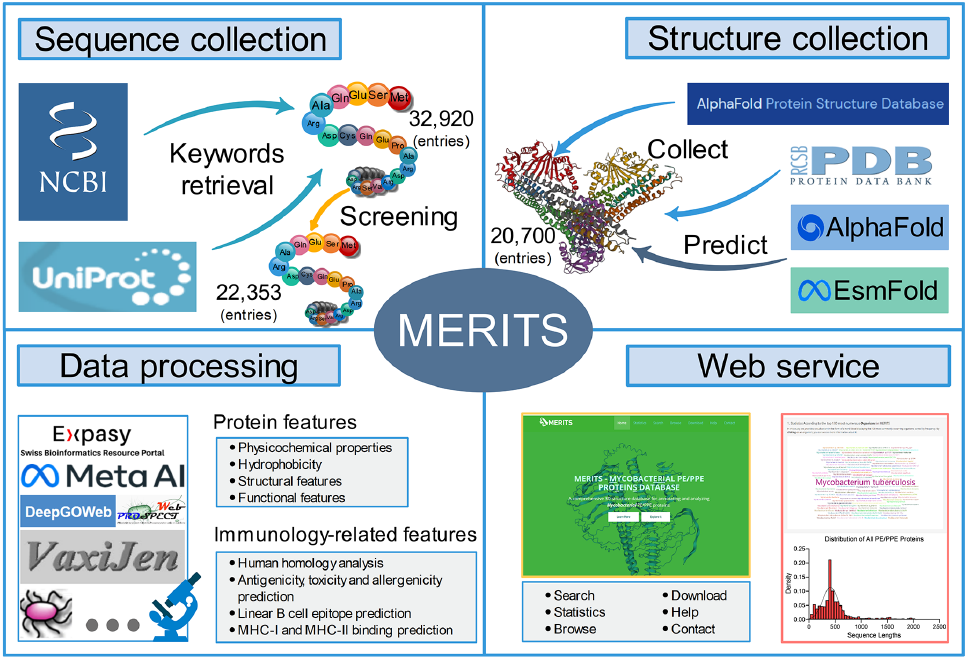
The overview of MERITS, including the collection of primary sequence and tertiary structure data for PE/PPE family proteins, sequence-structure-function annotation analysis of PE/PPE family proteins and database access.

In response to this critical gap, we have developed the MERITS (**M**ycobact**ERI**al PE/PPE pro**T**ein**S**) database. MERITS is a comprehensive *Mycobacterial* PE/PPE protein repository containing 22,353 non-redundant records curated from multiple sources. It offers extensive sequence-structure-function annotations, including physicochemical properties, subcellular localization, phosphorylation sites, and detailed protein function analysis. Additionally, it provides insights into immunology-related aspects such as antigenicity, toxicity, allergenicity, human homology, and epitope predictions. This wealth of information is pivotal for advancing our understanding of *Mycobacteria*, aiding in developing new anti-tuberculosis drugs, enhancing existing therapies, and creating innovative diagnostic methods for drug-resistant strains. By offering comprehensive and detailed data on PE/PPE proteins, MERITS paves the way for groundbreaking research and applications in combating *Mycobacterial* diseases.

## 2 Materials and Methods

### 2.1 Data collection

This section details the flowchart of the MERITS database as illustrated in **Figure1**. It encompasses the steps involved in collecting protein sequence and tertiary structure data for PE/PPE proteins, along with the subsequent sequence-structure-function annotation analysis. The specific methodologies and processes used in constructing the database and ensuring data accessibility are elaborated in the subsequent sections.

#### 2.1.1 Protein Sequence Data Collection

The initial step in constructing MERITS involved extracting protein data related to *Mycobacterium* PE/PPE proteins from the NCBI and UniProt databases. We used ‘*Mycobacterium*’ and ‘PE-PPE’ as search keywords, yielding 32,920 raw protein entries. To ensure the data’s relevance and quality, these entries underwent a rigorous validation and integration process based on specific criteria: (i) inclusion of proteins classified under the *Mycobacterium* PE/PPE protein family; (ii) confirmation that the ‘Taxonomy’ field indicates association with *Mycobacterium*; (iii) exclusion of entries with terms such as ‘partial’, ‘part’, ‘fragment’, ‘predicted’, ‘model’, ‘inferred’, ‘putative’, and ‘hypothetical’ in the Definition field; and (iv) avoidance of proteins from ‘uncultured’ organisms in the ‘Organism’ field. This meticulous selection process resulted in a curated collection of 22,353 entries, comprising the foundational data for *Mycobacterial* PE/PPE proteins in the MERITS database.

#### 2.1.2 Protein 3D Structure Data Collection

Tertiary structures of PE/PPE proteins are crucial for understanding *mycobacterial* growth, metabolism, and pathogenesis, and they aid in identifying new drug targets for antibiotic development. To collect accurate structures, we sourced experimentally verified PDB structures from RCSB PDB (Kouranov, 2006). Additionally, the predicted structures of AlphaFold (Jumper *et al*., 2021) were obtained from AlphaFold DB (Varadi *et al*., 2022). In instances where tertiary structures were unavailable in these databases, we utilised EsmFold (Lin *et al*., 2023), an efficient and nearly as accurate sequence-to-structure predictor as alignment-based methods, for generating predictions. Through these methods, we compiled a collection of 20,700 tertiary structures. However, it is important to note that PDB structure predictions for longer protein sequences were limited due to experimental constraints. The details of our tertiary structure data collection are summarised in **Table 1**.

**Table 1.**
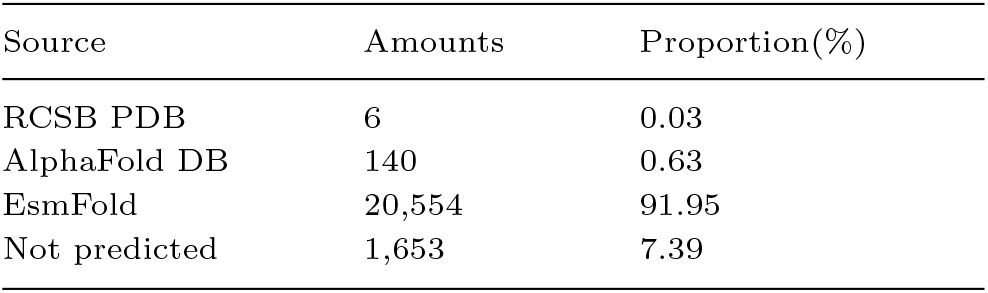
Results of tertiary structure data collection

| Source | Amounts | Proportion(%) |
| --- | --- | --- |
| RCSB PDB | 6 | 0.03 |
| AlphaFold DB | 140 | 0.63 |
| EsmFold | 20,554 | 91.95 |
| Not predicted | 1,653 | 7.39 |

### 2.2 Protein features calculation

The comprehensive analysis of protein features is fundamental for understanding the intricate biological roles of proteins. **Figure 2(C)** presents various panels illustrating the physicochemical properties, structural features, and functional attributes of proteins.

**Fig. 2.**
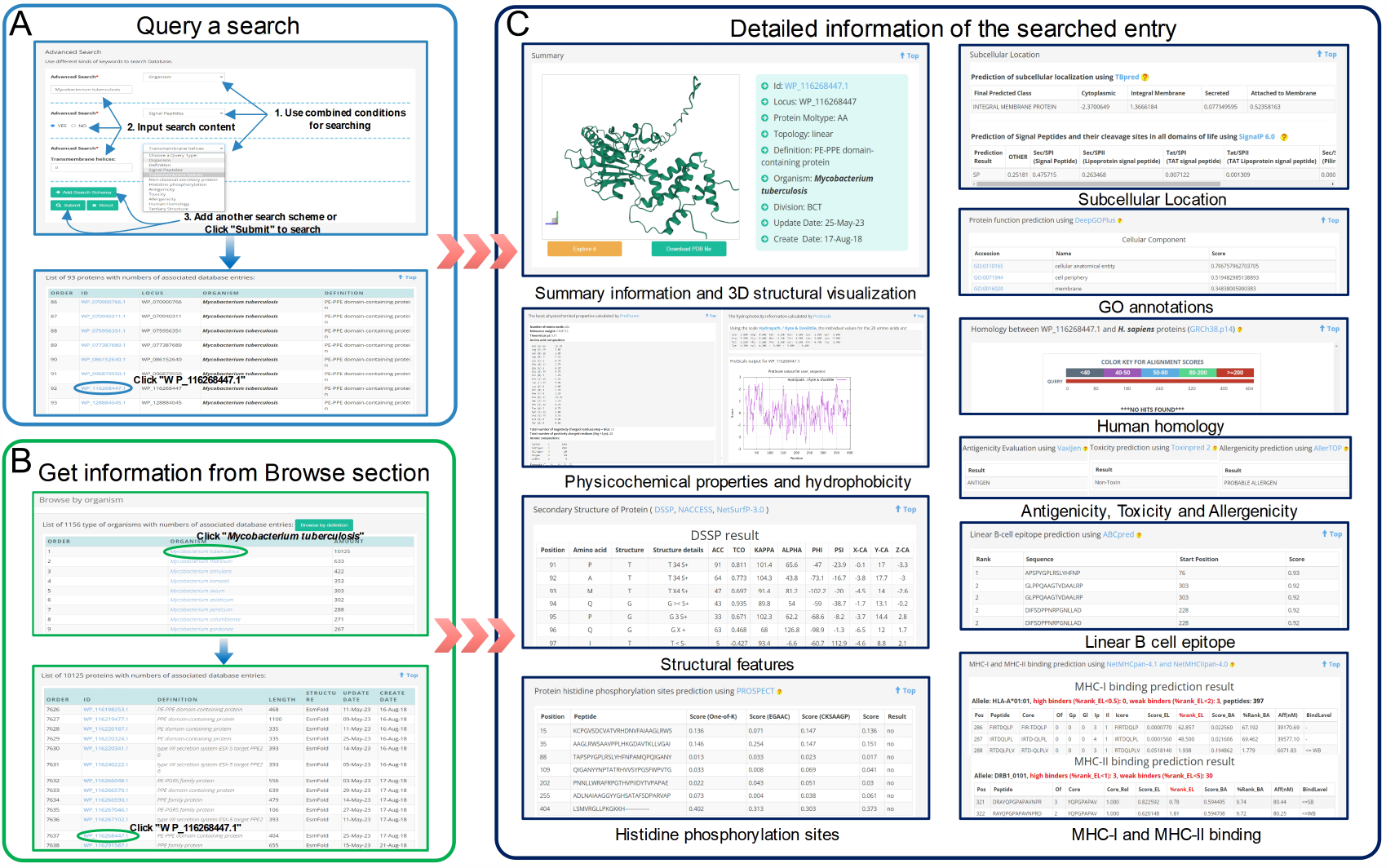
Two different methods for data accessing in MERITS. (A) An example showing how to use the advanced search function of MERITS. Here, three different search schemas were combined, Organism (*Mycobacterium tuberculosis*), Signal Peptides (YES), Transmembrane Helices (0); (B) An example illustrates how to explore the protein data through the Browse webpage (*Mycobacterium tuberculosis* and ID WP 116268447.1 is selected); (C) Typical search result webpage using the ID WP 116268447.1 as an example. The entry webpage consists of ten significant categories of information, comprising summary information and 3D structural visualization, Physicochemical properties and hydrophobicity, structural features, histidine phosphorylation sites, subcellular location, GO annotations, human homology, antigenicity, toxicity, allergenicity, linear B cell epitope, MHC-I and MHC-II binding.

#### 2.2.1 Physicochemical Properties and Hydrophobicity

Protein characteristics such as chemical composition, molecular weight, solubility, isoelectric point, amino acid composition, and spatial configuration significantly influence their structure, function, stability, and antigenicity. Hydrophobicity, dictated by amino acid composition, informs protein folding patterns. This study analysed these properties using ProtParam and ProtScale (Gasteiger *et al*., 2005).

#### 2.2.2 Structural Features

The structural integrity of proteins, vital for their functional analysis, was assessed using DSSP (Joosten *et al*., 2011; Kabsch and Sander, 1983) for secondary structure characteristics and NACCESS (Hubbard and Thornton, 1992) for atomic accessibility, as shown in the ‘Structural features’ panel of **Figure 2(C)**. Where 3D structures were unavailable, NetSurfP 3.0 (Høie *et al*., 2022) provided secondary structure predictions from amino acid sequences.

#### 2.2.3 Functional Features

##### (i) Protein Phosphorylation

Protein phosphorylation, specifically histidine phosphorylation, regulates cellular pathways (Li *et al*., 2020). PROSPECT (Chen *et al*., 2020) facilitated rapid and accurate histidine phosphorylation site predictions, depicted in the ‘Histidine phosphorylation sites’ panel of **Figure 2(C)**.

##### (ii) Subcellular location

Understanding the subcellular localisation of proteins is integral to its functionality (Wang *et al*., 2022). TBpred (Rashid *et al*., 2007) classified *Mycobacterial* proteins into cytoplasmic, integral membrane, secretory, or membrane-attached by lipid anchor categories. Signal peptides and transmembrane helices, indicative of secretory proteins, were predicted with SignalP 6.0 (Teufel *et al*., 2022) and TMHMM 2.0 (Krogh *et al*., 2001). Non-classical secretion pathways were analysed using SecretomeP-2.0 (Bendtsen *et al*., 2004), as illustrated in the ‘Subcellular location’ panel of **Figure 2(C)**.

##### (iii) Gene Ontology (GO) Annotation

GO annotations, essential for linking proteins to disease pathology and drug discovery, were generated using DeepGOPlus (Kulmanov and Hoehndorf, 2020), a deep learning approach that combines sequence-based predictions with similarity-based predictions for Cellular Components, Molecular Functions, and Biological Processes. These are displayed in the ‘GO annotations’ panel of **Figure 2(C)**.

### 2.2 Immunology-related features

#### 2.3.1 Human homology analysis

For vaccine candidacy, non-homologous human proteins are considered potential targets (Mohinani *et al*., 2021). We sourced human protein data from Homo sapiens GRCh38.p13 of the NCBI database and used BLASTp (E-value set to 0.05) to compare homology between PE/PPE proteins and H. sapiens proteins, assessing their suitability as vaccine candidates. **Figure 2(C)** includes an example in the ‘Human homology’ panel.

#### 2.3.2 Antigenicity, toxicity and allergenicity prediction

The comprehensive immunological profile of PE/PPE proteins, encompassing antigenicity, toxicity, and allergenicity, was analysed using VaxiJen 2.0 server (Doytchinova and Flower, 2007), ToxinPred 2 (Sharma *et al*., 2022), and AllerTop 2.0 server (Dimitrov *et al*., 2014). These tools predict immunological responses based on protein sequences, with results illustrated in **Figure 2(C)** under the ‘Antigenicity, Toxicity, and Allergenicity’ panel.

#### 2.3.3 Linear B-cell epitope prediction

Identifying B-cell epitopes is essential for vaccine design, diagnostics, and allergy research. ABCpred (Saha and Raghava, 2006), which uses recurrent neural networks for accurate linear B-cell epitope prediction, provided results, including ranking and scores, shown in the ‘Linear B cell epitope’ panel of **Figure 2(C)**.

#### 2.3.4 MHC-I and MHC-ll binding prediction

The binding affinity of peptides to MHC molecules is critical for cellular immunity and is predictive of immunogenicity. Using NetMHCpan-4.1 and NetMHCIIpan-4.0 (Reynisson *et al*., 2020), we predicted peptide binding to MHC-I and MHC-II, selecting alleles covering major human ethnic groups for MHC-I (A1, A2, A3, A24, B7) and HLA-DR, HLA-DQ, HLA-DP, and H-2-1 for MHC-II. **Figure 2(C)**, the ‘MHC-I and MHC-II binding’ panel, presents predictive results and examples of MHC binding.

### 2.4 Database Access and Navigation

An intuitive and powerful access interface is critical for the practical utility of any comprehensive database. MERITS addresses this need with a suite of user-friendly functionalities designed for streamlined navigation and data retrieval. The “Home page” of MERITS serves as the gateway, providing a succinct introduction to the database’s capabilities and guiding users on how to leverage its resources. This initial interface is designed to familiarise users quickly with the overall structure and functionalities of MERITS. On the “Statistics page”, users are presented with visual statistical analyses that elucidate the scope and distribution of the PE/PPE protein data within MERITS. These visualisations help convey the depth and breadth of the collected data, offering insights into the diversity of the proteins catalogued. For targeted data inquiry, the “Search page” is equipped with a versatile search engine that supports multiple query methods. Users can perform an ID search for direct retrieval of specific entries, a keyword search for broader data exploration, or utilise the advanced search function. The advanced search capability is particularly robust, enabling the combination of various criteria such as organism type, protein definition, presence of signal peptides or transmembrane helices, antigenicity, human homology, and available tertiary structures. This feature is indispensable for researchers requiring highly specified data sets. The “Browse page” allows for exploratory navigation, offering users the opportunity to peruse the database’s content systematically. It is structured to facilitate browsing by definition or organism, providing a comprehensive overview of the PE/PPE proteins catalogue. Lastly, the “Download page” caters to offline data analysis needs. Users can download various data formats, including SQL files for database integration, files detailing structural features for computational studies, and datasets of histidine phosphorylation sites pertinent to functional analysis. This functionality underscores MERITS’s commitment to open science by enabling unrestricted access to its data repository for further research and development. Collectively, these features ensure that MERITS is not just a repository of information, but a dynamic tool that empowers researchers to conduct effective and informed scientific inquiries into the *Mycobacterium* genus.

### 2.5 Implementation and Visualization

MERITS is underpinned by a robust MySQL relational database chosen for its efficiency in handling complex data relationships. The server-side scripting is powered by PHP, while Python scripts handle data processing tasks, ensuring a seamless back-end operation. The user interface, crafted with user experience as a priority, employs the jQuery framework for dynamic content manipulation, enhancing interactivity and responsiveness. The web interfaces’ fundamental operations rely on JavaScript and jQuery, providing a smooth and interactive user experience. The visual aesthetic and structural layout of the pages are designed using the Bootstrap framework, known for its responsive design templates. The entire platform operates on a dedicated Linux server, equipped with a 4-core CPU and 16 GB of memory, providing the necessary computational power and storage capacity, with a 200 GB hard disk, to handle substantial data processing tasks. Nginx is the chosen web server for its high performance and stability.

On the statistics page, MERITS provides five distinctive statistical analyses, reflecting the multidimensional nature of the data. Interactive pie charts, crafted with ECharts (Li *et al*., 2018), offer users an engaging way to comprehend the data, where hovering a mouse over chart segments reveals deeper insights, fostering an exploratory data experience. The entry page is enriched with 3D protein structural visualisations powered by PDBe (Nair *et al*., 2021). This integration allows users to interact with complex molecular structures directly within their web browser without the need for additional plugins or Java, thanks to PDBe’s dual functionality as a JavaScript plugin and a web component. For more in-depth sequence analysis, we incorporated BlasterJS (Blanco-Míguez *et al*., 2018) to enable the intuitive display of sequence alignments, enhancing the understanding of homology and sequence conservation. These sophisticated visualisation tools, exemplified in the ‘Summary information and 3D structural visualisation’ and ‘Human homology’ panels of **Figure 2(C)**, are pivotal in making MERITS not just a database but an interactive platform that brings molecular data to life, allowing researchers to visualise, analyse, and interpret biological data within a comprehensive and user-centric environment.

## 3 Results and Discussion

### 3.1 Enhanced Search Functionalities in MERITS

MERITS provides multiple search functionalities, allowing users to access data directly and comprehensively. This section elaborates on these search capabilities with a detailed description and demonstrations of MERITS’s features.

The platform supports three primary search strategies. The ‘ID search’ enables data retrieval via specific Protein ID (e.g. ‘WP 116268444.1’) or gene locus tags (e.g. ‘PPE54 MYCTU’). For broader inquiries, the ‘keyword search’ allows entries by based on organism names or protein definitions. Both methods offer “Example” buttons — intuitive guides that populate search fields with sample data, thereby simplifying the user experience. The ‘advanced search’ functionality stands out with its comprehensive query options. It boasts eleven different search patterns, catering to various research needs. Users can specify parameters such as organism type, protein definition, presence of signal peptides, number of transmembrane helices, non-classical secretion, histidine phosphorylation presence, and antigenicity level. This feature is enhanced by a user-friendly interface where the ‘Add Search Schema’ button enables complex, multi-criteria searches for nuanced data mining. An illustrative search example in **Figure 2(A)** showcases how a user can filter for secreted proteins from ‘*Mycobacterium tuberculosis*’, marked by signal peptides and the absence of transmembrane helices. The resulting data is displayed in a detailed table format, accessible via a dedicated results webpage, as depicted in **Figure 2(A)**. This table is organised into five critical columns that encapsulate the essence of the search query, offering at-a-glance information on ID, locus, organism, and protein definition.

For an in-depth exploration, each Protein ID in the results table is a hyperlink to a detailed annotation page. Clicking on these links unveils additional information, such as 3D structural visualisations, detailed physicochemical profiles, structural & functional features, immunological properties, and potential epitopes. These annotations are crucial for researchers and are visually summarised in the ‘Summary information and 3D structural visualisation’ and ‘Human homology’ panels of **Figure 2(C)**. MERITS thus not only aggregates an extensive dataset but also furnishes the field with sophisticated search tools. These tools are designed to streamline data retrieval, encouraging the exploration of PE/PPE proteins’ multifaceted roles and fostering the development of potential diagnostic and therapeutic applications.

### 3.2 Database contents

This section provides an in-depth statistical analysis of the PE/PPE proteins within MERITS, helping to contextualise the database’s content. **Figure 3(A)** details the amino acid composition distribution among PE/PPE proteins. It reveals a significant clustering of protein lengths, with the most common range being 350 to 400 amino acids, accounting for 15.8% of the dataset. Notably, the data indicates that over 90% of the PE/PPE proteins are less than 1000 amino acids in length, suggesting a concentration in sequence sizes and potentially implicating functional or structural constraints in their evolution. The amino acid frequency distribution, shown in **Figure 3(B)**, identifies Glycine (G) as the most abundant residue, comprising 31.68% of the amino acids in PE/PPE proteins. This preponderance aligns with findings from previous studies [36, 37], hinting at the importance of Glycine in the structural formation of these proteins. Organism distribution within the MERITS database is graphed in **Figure 3(C)**, with M. tuberculosis emerging as the predominant species. It is followed by *M. marinum, M. simulans, M. kansasii*, and *M. avium*, with these top five species representing about 80% of the organismal data, reflecting the research focus on certain *mycobacterial* species due to their clinical significance. In **Figure 3(D)**, the distribution of different PE/PPE protein families is analysed. The PE-PGRS family protein emerges as the most frequently occurring, constituting 30.7% of the dataset, followed by proteins containing the PE domain at 20.9%. This dominance suggests these families play substantial roles in the genus’s pathogenic mechanisms or immune interactions.

**Fig. 3.**
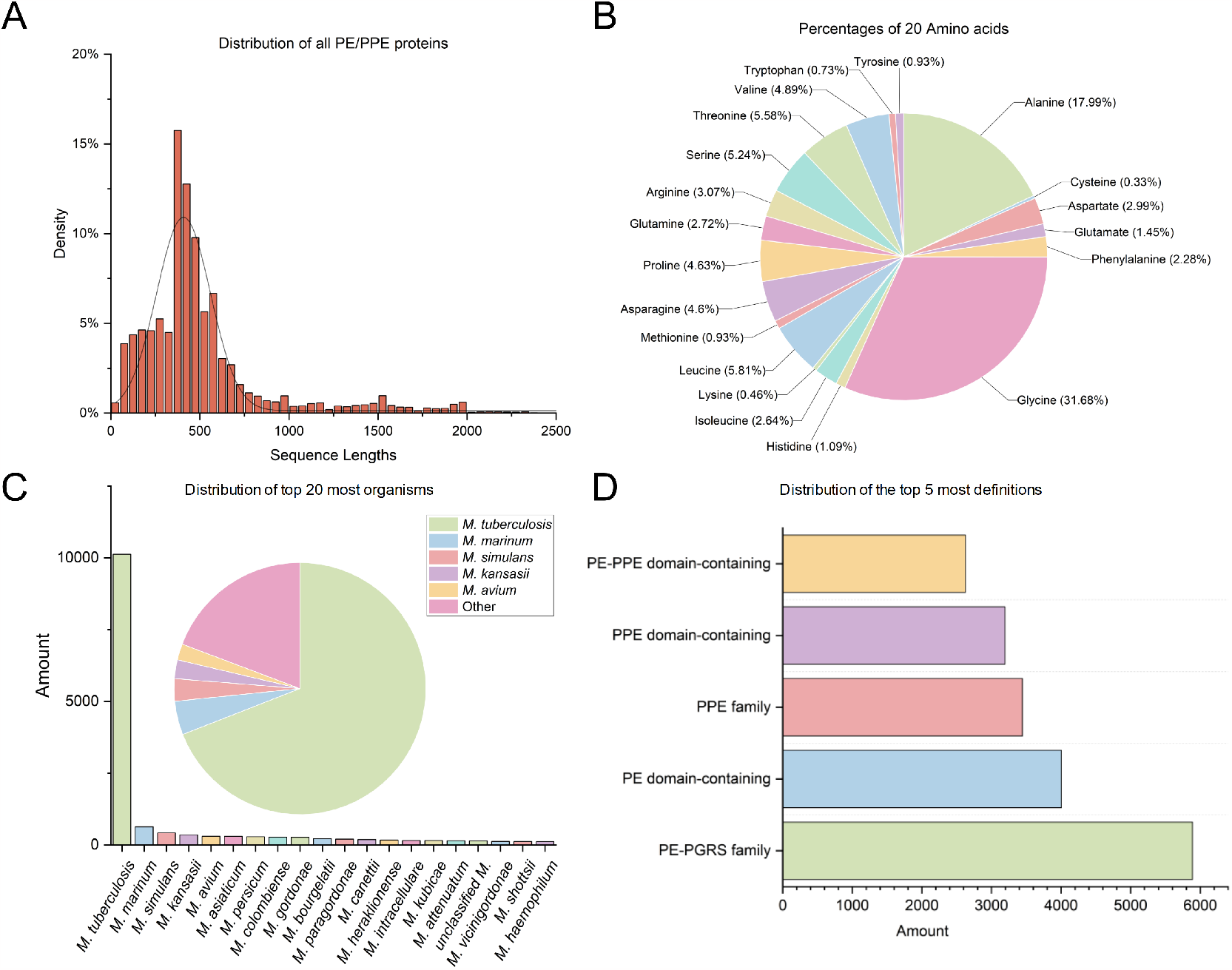
Statistical analysis results of PE/PPE proteins. (A) Distribution of all PE/PPE proteins according to their protein sequence lengths; (B) Frequency distributions of 20 amino acids in all accumulated PE/PPE proteins; (C) Distribution of top 20 most organisms; (D) Distribution of the top 5 most definitions.

The secondary structure distribution of these proteins is depicted in **Figure 4(A)**. The *α*-helix (H) structure is overwhelmingly the most common, at 59.91%, while the *π*-helix (I) is rare, forming a mere 0.58% of the dataset. Such prevalence of *α*-helices may indicate structural stability or specific functional interactions in PE/PPE proteins. Subcellular localisation, crucial for understanding protein function, is visualised in **Figure 4(B)**. TBpred analysis shows that most (77.99%) of PE/PPE proteins are ‘Integral Membrane Proteins’, underscoring the importance of membrane association in their biological roles. Lastly, **Figures 4(C)** and **(D)** present the predicted distribution of cytotoxic T-lymphocyte (CTL) and helper T-lymphocyte (HTL) epitopes. NetMHCpan predicts a total of 232,908 CTL epitopes, with a substantial number of strong binders to common alleles such as A1, A2, A3, A24, and B7. In contrast, relevant for HTL responses, MHC II binders are most abundant for H-2-1 alleles, with 1,591,814 predicted epitopes, highlighting their potential for vaccine development. These comprehensive statistical insights not only delineate the breadth of data within MERITS but also underscore the extensive biological significance of PE/PPE proteins, paving the way for future research endeavours into vaccine development and disease understanding.

**Fig. 4.**
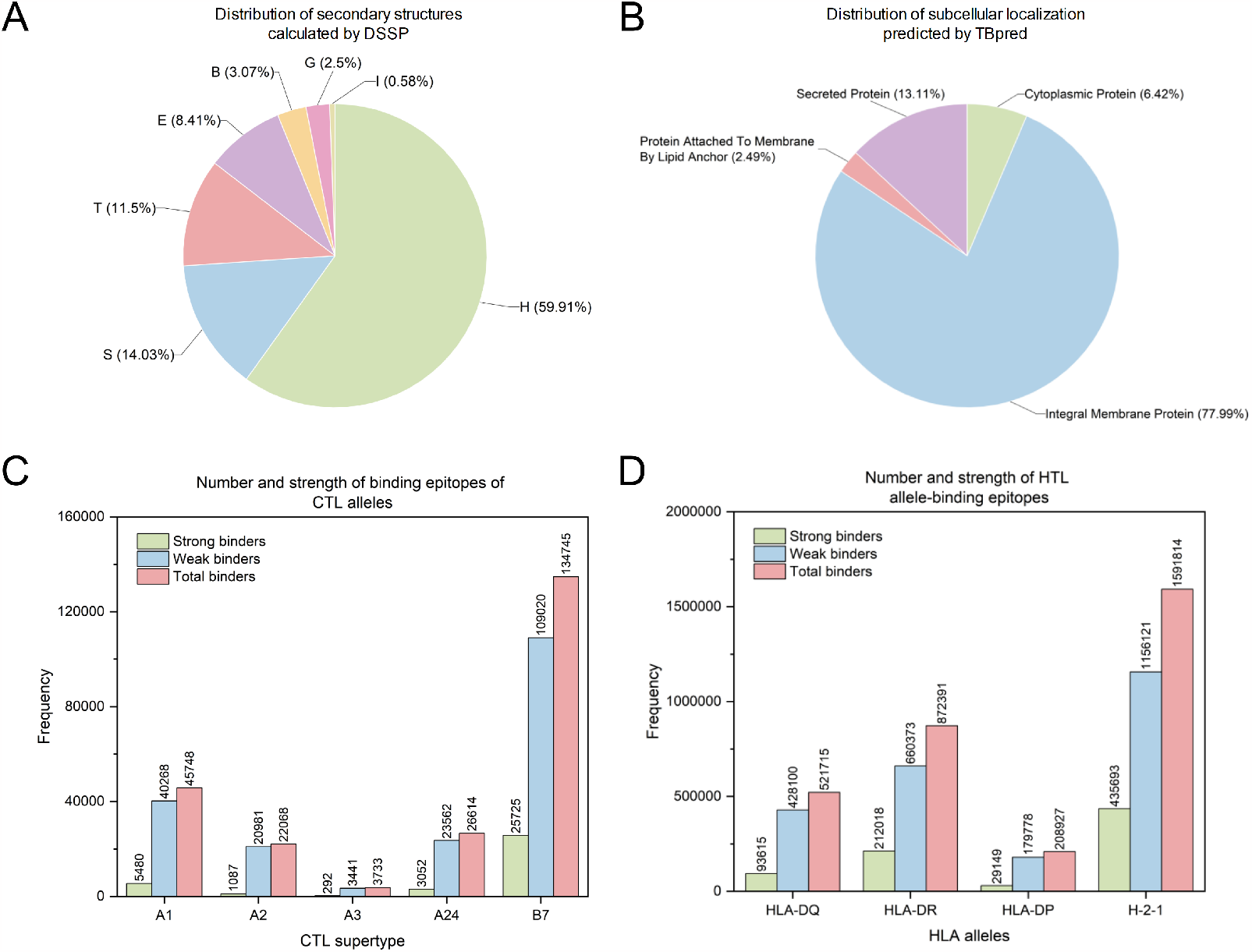
Part of structural and functional features of PE/PPE proteins. (A) Distribution of secondary structures calculated by DSSP; (B) Distribution of subcellular localization predicted by TBpred. (C) Number and strength of binding epitopes of CTL alleles predicted by NetMHCpan-4.1. (D) Number and strength of HTL allele-binding epitopes predicted by NetMHCIIpan-4.0.

## 4 Conclusions

This study proposed MERITS, which is a pivotal and freely accessible database dedicated to the comprehensive analysis of *Mycobacterial* PE/PPE proteins, vital for understanding the *Mycobacterium* genus. It aggregates a wealth of sequence and structural data from various sources, meticulously analysing sequence-structure-function relationships. This integration offers insights crucial for the fields of bioinformatics, vaccine development, and *Mycobacterial* research. The platform excels in visualising 3D structures and delineating the structural and functional attributes of a vast array of PE/PPE proteins. It also enriches our understanding of immunological aspects by annotating antigenicity, toxicity, and MHC-I/MHC-II binding affinities. MERITS acts not just as a repository but as an analytical tool, deepening our grasp of PE/PPE protein mechanisms and functionalities. Engineered with PHP and JavaScript, MERITS offers an intuitive interface for efficient data retrieval and interactive visualisation, employing bioinformatics tools like PDBe and BlasterJS for an enhanced user experience. The database is designed to be dynamic, with biannual updates ensuring its relevance and accuracy.

Looking forward, MERITS aims to expand its scope to include protein-protein interaction networks, recognising their fundamental role in cellular processes and disease mechanisms (Szklarczyk *et al*., 2021). This addition will further augment the database’s research utility. Moreover, we anticipate leveraging advancements in large language models (LLMs) to enrich the database’s analytical capabilities. This integration could revolutionise the analysis of PE/PPE proteins, particularly in sequence-structure-function correlations, offering a more nuanced understanding and facilitating rapid hypothesis generation and testing.

As the premier comprehensive database for *Mycobacterial* PE/PPE proteins, MERITS is positioned to be an invaluable asset in the scientific community. It promises to catalyse significant advancements in understanding *Mycobacterium*, aiding in developing novel therapeutics and vaccines, and enhancing our overall comprehension of these critical biological entities.

## Acknowledgments

This work is supported by the National Natural Scientific Foundation of China (No. 62202388), the National Key Research and Development Program of China (No. 2022YFF100 0100), the Qin Chuangyuan Innovation and Entrepreneurship Talent Project (No. QCYRCXM-2022-230), Talent Research Funding at Northwest A&F University (No. Z1090222021) and the Major and Seed Inter-Disciplinary Research Projects awarded by Monash University.

## Data availability

The data of this study is available at http://merits.unimelb-biotools.cloud.edu.au/index.php/download

